# Preferred left-handed conformations of glycyls with pathogenic variants protect against aggregation

**DOI:** 10.1101/2024.02.09.579738

**Authors:** Purva Mishra, Rajesh Potlia, Kuljeet Singh Sandhu

**Affiliations:** DBT Bioinformatics Centre; Department of Biological Sciences; Indian Institute of Science Education and Research (IISER) – Mohali Sector 81, SAS Nagar 140306, India

## Abstract

Due to the lack of C***β*** atom, the glycyl residues can exhibit left-handed Ramachandran conformations that are mostly disallowed for L-amino acids. The structural and functional significance of distinct glycyl conformations remains under-appreciated. Through systematic analyses of various datasets, we show that: i) the left-handed glycyl residues are over-represented at disease-associated sites and are evolutionarily conserved. ii) The mutations of L-disallowed Gly tend to destabilize the native folding as assessed through the change in free energies. An independent analysis of folding nuclei further corroborates the findings. iii) L-disallowed Gly are enriched at the aggregation gatekeepers, more significantly so in thermophiles, and the mutations thereof reduce the protein solubility. (iv) The positive ***Φ*** dihedral angle of L-disallowed Gly disorients its C***α*** atom out of the phase of alternating pleats of ***β***-strand, conforming a crescent that is incompatible to further pair with other ***β***-strands, and thus discourages the inter-molecular aggregation of β-strands during protein folding. v) L-disallowed conformation of Gly holds predictive power to identify sites having pathogenic variants. Altogether, our observations highlight that the L-disallowed conformations of glycyls are evolutionarily selected to endow protein stability and protection against aggregation. Apart from enhancing the existing knowledge, the findings have implications in prioritizing the genetic lesions implicated in diseases, and in designing proteins with greater stability and solubility.

## Introduction

Genome-wide association studies are instrumental in addressing the disease associations of single nucleotide polymorphisms (SNPs). Thousands of disease-associated SNPs map to protein-coding regions and are missense in nature^1^. The pathogenic missense variants alter a distinct set of functional pathways than the benign ones, causing the disease phenotypes^2^. A major challenge has been to understand how missense mutations alter the structural and functional properties of the proteins, leading to perturbations of the pathways implicated in diseases. Yue *et al.* has shown that a significant proportion of pathogenic missense variants are buried within protein structure and that the conformational destabilization is the leading cause of deleterious effect of missense variants^3^. More recently, a whole repertoire of disease-associated protein structural features was uncovered, which included changes in posttranslational modification sites, physicochemical properties of residues, residue exposure level, etc., apart from functional interaction sites^4^. Several algorithms have also been developed to predict the deleterious impact of missense variants^5–15^. Attempts to delineate the conformational features associated with the disease-associated mutations though remains limited.

Ramachandran plot of ***Φ*** and ***Ψ*** dihedral angles is fundamental to the conformational analysis of proteins and peptides^16^. The ***Φ****-****Ψ*** torsions for the non-proline and non-glycyl L-amino acid residues are generally constrained within three designated areas in the Ramachandran plot: the extended ***β-***sheet region (E), the right-handed ***α***-helical region (R), and the left-handed ***α-***helical region (L) due to various steric clashes^16^. Unlike other residues, glycyl residues exhibit greater torsional freedom due to the lack of C***β*** atom^17^. The major difference lies towards positive ***Φ*** values. Apart from the L-region, the right-hand side of the Ramachandran plot is mostly disallowed, referred as L-disallowed, for non-glycyl residues, but is allowed for Gly. This may have some important implications for the structure and function of the proteins since any mutation of the L-disallowed Gly to non-Gly L-amino acid may require significant alteration in the backbone conformation. Decades ago, an analysis of lysozyme had revealed that the mutations of Gly exhibiting Ramachandran conformation near left handed helical region leads to relatively destabilizing free energies^18^. The glycyl residues having left-handed helical conformation are known helix stoppers and ***β***-turn makers^19–22^. Valiyaveetil *et al.* showed that a glycyl residue in potassium ion-channel serves as a surrogate to D-amino acid^23^. Mutation of Gly to L-Ala, but not to D-Ala, leads to loss of structural and functional attributes of protein^23^. Prakash *et al.*^24^ has shown that the glycyl residues occurring in the evolutionarily invariant peptides in bacteria tend to be buried and were moderately more frequent in the L-disallowed regions. More recently, Lakshmi *et al.* has claimed that the small size of the Gly, regardless of L-allowed or L-disallowed conformation, could likely have guided their evolutionary selection in conserved protein families^25^. The underlying significance of L-disallowed conformation of glycyl residues, therefore, remains enigmatic.

In the present study, we attempted to understand the implication of L-disallowed conformation of Gly in human diseases and the mechanistic basis thereof. Through systematic analysis, we showed a significant representation of L-disallowed Gly at disease-associated and aggregation gatekeeping sites. The mutations of L-disallowed Gly to non-Gly residues destabilize the protein conformation, reduces protein solubility, and promotes aggregation. The L-disallowed conformation of Gly disrupts the atypical alternating pattern of ***β***-pleats, disfavoring the further paring of beta-strands and consequently reducing aggregate forming potential. Our observations also had predictive power to improve the inference of missense variants.

## Results

### Glycyls having pathogenic variants frequently exhibit left-handed conformation

Analysis of 1104 pathogenic and 343 benign variants of Gly from the ADDRESS database revealed that the Gly associated with the pathogenic variants more often exhibited left-handed conformation (+ve ***Φ***) when compared with the ones associated with benign variants (Figure 1A-D, Table S1). The observation was consistent for buried and surface-exposed residues with some differences (Odd-ratio_surface_ =1.37, 95% lower CI: 1.019; P_Fisher, surface_=0.04, Odd-ratio_buried_ =1.32, 95% lower CI: 0.973, P_Fisher, buried_ =0.07, Figure 1A-D). Motivated by this observation, we further categorized glycyl residues into L-allowed and L-disallowed categories using arbitrary ***Φ-Ψ*** range covering allowed regions in the Ramachandran plot of L-amino acids for detailed analysis (Figure S1). We confirmed that the glycyls at the pathogenic sites tend to exhibit L-disallowed conformation more often than those at the benign sites (Odd-ratio_surface_ =1.5, P_Fisher, surface_=0.01, Odd-ratio_buried_=1.32, P_Fisher, buried_=0.07).

**Figure 1.**
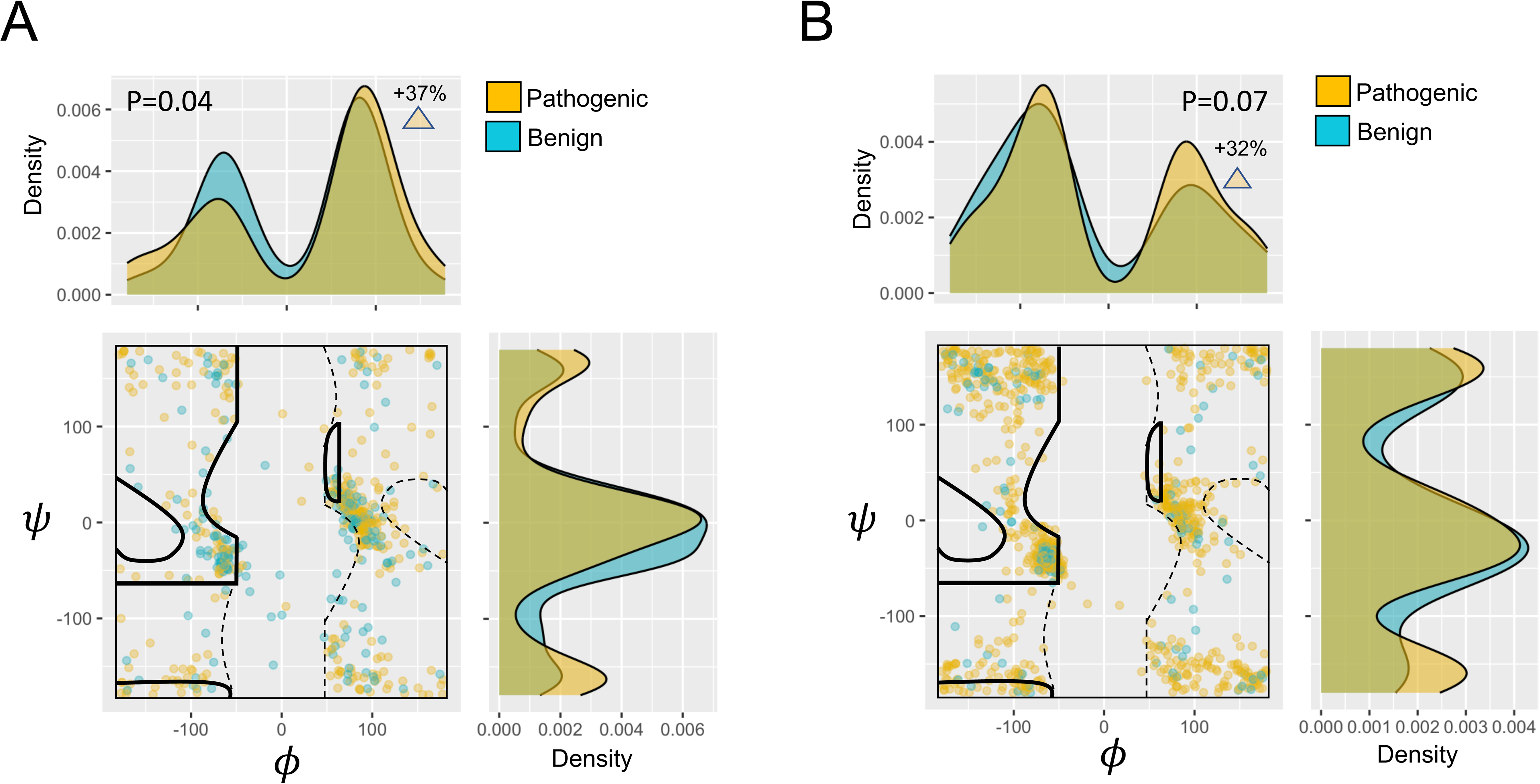
Conformation of glycyl residues having pathogenic or benign variants. Ramachandran plots of (A) surface exposed and (B) buried glycyl residues at benign and pathogenic sites. The individual density plots of " and # are aligned accordingly. P-values were calculated for +ve and -ve ***Φ*** values of Gly at pathogenic (golden) and benign (aquamarine) sites using one-sided Fisher’s exact tests. The percentage shown represents the increase in probability of observing left-handed gly conformation over right-handed conformation at pathogenic sites when compared to that at benign sites.

### L-disallowed glycyls are evolutionarily more conserved than L-allowed ones

Since the pathogenicity of variants generally correlates with the evolutionary conservation of the underlying residue, we compared the residue-wise evolutionary rates for the L-allowed and L-disallowed Gly at the pathogenic and the benign sites. The analysis showed significantly lower evolutionary divergence rates of L-disallowed Gly as compared to L-allowed ones at both pathogenic and benign sites, though relatively greater at pathogenic sites (Figure 2A, Table S1). This was consistent with the earlier reports wherein the Gly present in the conserved regions of proteins tended to exhibit L-disallowed conformation^24^.

**Figure 2.**
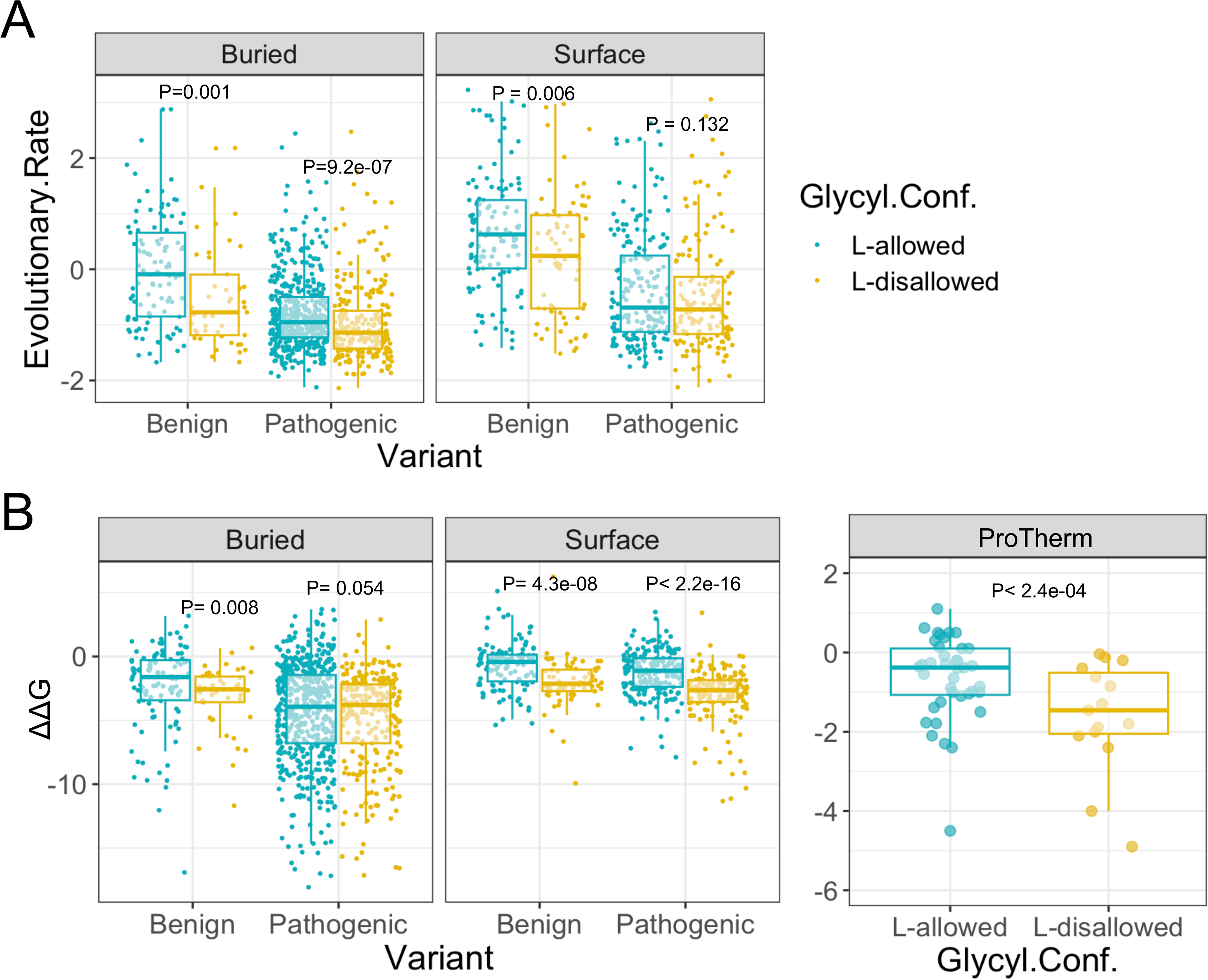
Evolutionary rates and the Gibbs free energy changes (upon mutation) for L-allowed and L-disallowed glycyl residues. (A) Residue-wise evolutionary rates of surface-exposed and buried glycyls with L-allowed and L-disallowed conformations. (B) Gibbs free energy changes (ΔΔG = ΔG_variant_ - ΔG_WT_) for surface exposed and buried glycyl residue with L-allowed and L-disallowed conformations. We obtained the ΔΔG values from FoldX and inversed the signs (Methods section). (C) Experimentally estimated Gibbs free energy changes for L-allowed and L-disallowed glycyl residues. We calculated the P-values using Mann-Whitney U tests.

### L-disallowed glycyls implicate in the stability of native protein structures

To understand the role of L-disallowed residues, we first scanned their presence at the functional sites of the proteins. Out of total 406 L-disallowed Gly at pathogenic sites, only 2 mapped to DNA binding sites and none mapped to ligand, metal, and RNA binding sites (Table S2). The localization within functional sites did not, therefore, explain the greater occurrence of L-disallowed Gly at the pathogenic sites.

We then tested the hypothesis whether the L-disallowed conformation of Gly was selected to endow conformational stability to the proteins. We compared the ΔΔG values, obtained from the FoldX framework, for the Gly at the pathogenic and benign sites. Interestingly, mutations of L-disallowed Gly exhibited conformational destabilizing negative ΔΔG when compared with the L-allowed ones (Figure 2B, Table S1). We further scrutinized this observation by analyzing experimentally calculated ΔΔG values from the ProTherm database (Table S3). The results in Figure 2C consistently supported our claim. To test whether the observed impact of mutations on protein stability was not due to the charge and other properties of the mutated residue, we obtained all the disease-associated Gly-to-Ala mutations and re-assessed the ΔΔG values for this subset (Figure S2, Table S4). The results were similar to those observed for the whole dataset, suggesting that the role of L-disallowed conformation of Gly in protein stability cannot be dismissed by the arguments of side-chain properties, other than that of Ala, of the mutated residues.

### Glycyls at aggregation gatekeeper sites exhibit L-disallowed conformation

The partial unfolding or destabilization of protein structures can expose the aggregation-prone regions (APRs), leading to intermolecular aggregation^26^. APRs are generally rich in hydrophobic ***β***-sheet forming residues and are flanked by short stretches (1-3 residues) of aggregation gatekeepers (GKs)^26^. We observed significantly higher enrichment of ***β***-sheet residues at and adjacent to the disease-associated and the benign L-disallowed Gly when compared to those adjacent to L-allowed Gly (Figure 3A, Table S5). Instead, the L-allowed Gly were generally flanked by ***α***-helical residues (Figure 3A). The observation hinted that the L-disallowed Gly may serve as GKs. The GKs are generally enriched with charged (R, K, D, E) and hydrophilic amino acids (S, T, N) and, to a lesser extent, small amino acids (G, P)^26^. We, therefore, tested the amino acid composition of -1 upstream and +1 downstream of Gly at pathogenic and benign sites. We observed enrichment of charged (primarily Arg) and hydrophilic amino acids (Ser and His) up and downstream of disease-associated L-disallowed Gly when compared those of L-allowed ones, hinting that the L-disallowed Gly might be the part of GKs (Figure 3B, Table S6). There were a few hydrophobic amino acids (like Val) too flanking the disease-associated L-disallowed Gly, possibly marking the Gly located towards the end of GK and adjacent to the potential ***β*** -sheet forming region or the APR. The analyses of inferred APRs further showed that the aggregate-forming potential was higher in the adjacent regions but dropped significantly at L-disallowed Gly, possibly marking gate-keeper regions (Figure 3C, Table S7). The aggregate-forming potential around L-allowed Gly did not exhibit any such pattern though (Figure 3C). These observations coherently suggested that the L-disallowed Gly may gate-keep the APRs.

**Figure 3.**
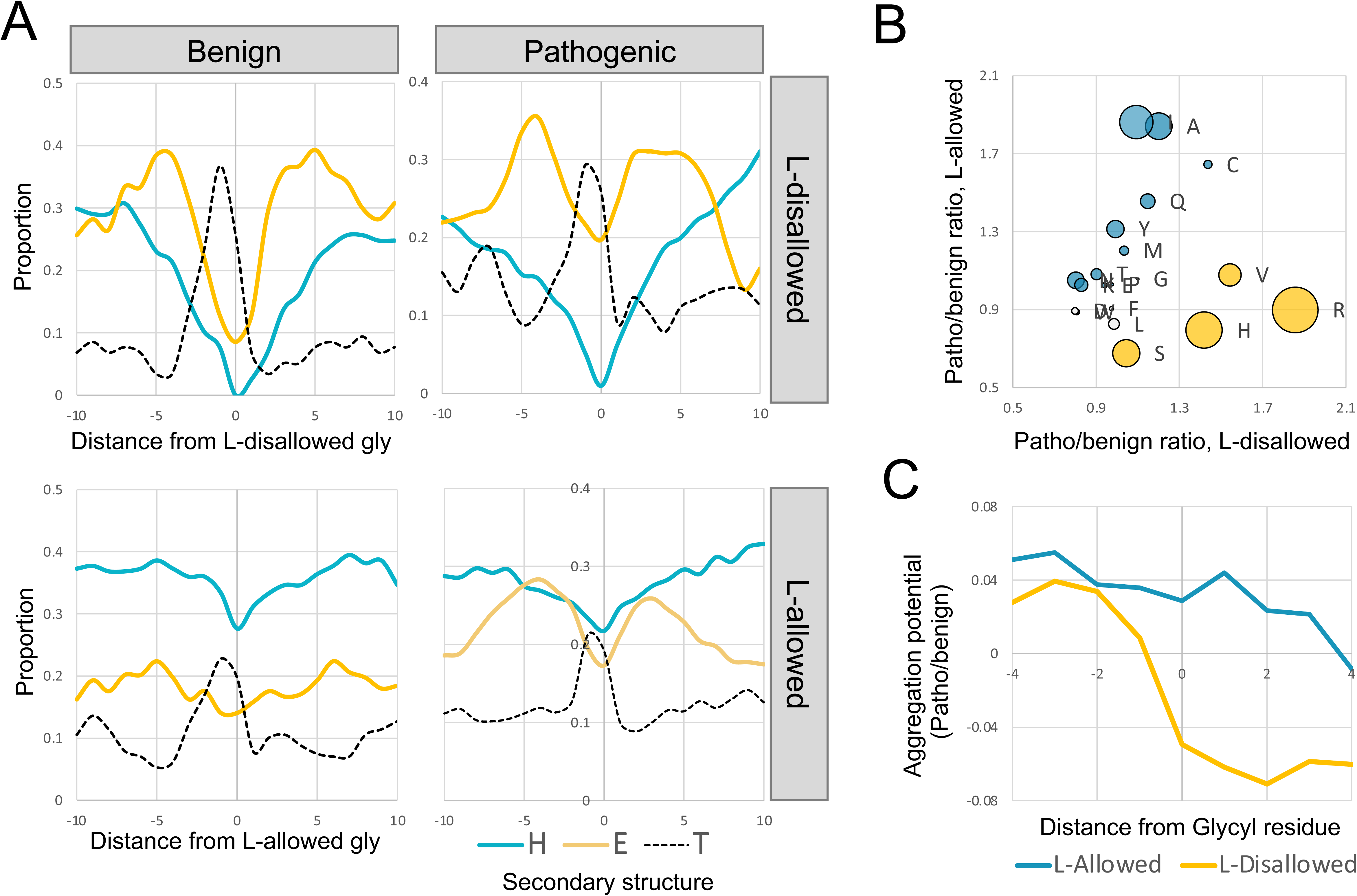
Aggregation related attributes around glycyl residues. (A) Mean enrichment of secondary structure states around L-disallowed and L-allowed glycyl residues at benign and pathogenic sites. (B) Bubble-plot of the amino acid frequencies (pathogenic-to-benign ratio) around L-disallowed glycyls (x-axis) and around L-allowed glycyls (y-axis). The bubble size signifies the log10(patho/benign) ratio. The aquamarine color marks the enrichment around L-allowed Gly, while the orange color highlights the enrichment around L-disallowed Gly. (C) The ratio of aggregation potential (pathogenic-to-benign) around L-allowed and L-disallowed Gly. We calculated the p-values using t-tests.

To further scrutinize the observed association of L-disallowed conformation of Gly with the aggregation gatekeeping, we obtained the APRs and the adjacent GKs for 373 mesophilic and 373 thermophilic homologous proteins for which structural coordinates were available^27^. Conformational analysis of Gly residues in APRs and GKs revealed a significantly greater representation of L-disallowed Gly in GKs when compared to APRs (Figure 4A, P=1.5e-04, Table S8). Interestingly, the observed association was statistically stronger for the thermophilic proteins when compared to mesophilic ones despite being homologous (Figure 4A, P=2.0e-06, Table S8). Unlike APRs, which are rich in hydrophobic amino acids and are mostly buried, GKs have hydrophilic/charged amino acids and may tend to be surface-exposed. This may create bias in our observations. We, therefore, controlled our analysis by only taking the buried GKs alongside buried APRs. The analyses consistently supported the L-disallowed conformation of glycyl residues in GKs (Figure S3, P=0.01 & 4.0e-03). Not only supporting our claim that the L-disallowed Gly tends to gate-keep the APRs, but the observation also implied the heightened selection of L-disallowed Gly at GK sites in thermophiles. We further obtained the glycyl mutations that were experimentally inferred to impact the solubility of proteins^28^. The Gly associated with decrease in protein solubility upon mutation mostly exhibited L-disallowed conformation when compared with the ones that were associated with increased solubility or were neutral (Figure 4B, Table S9).

**Figure 4.**
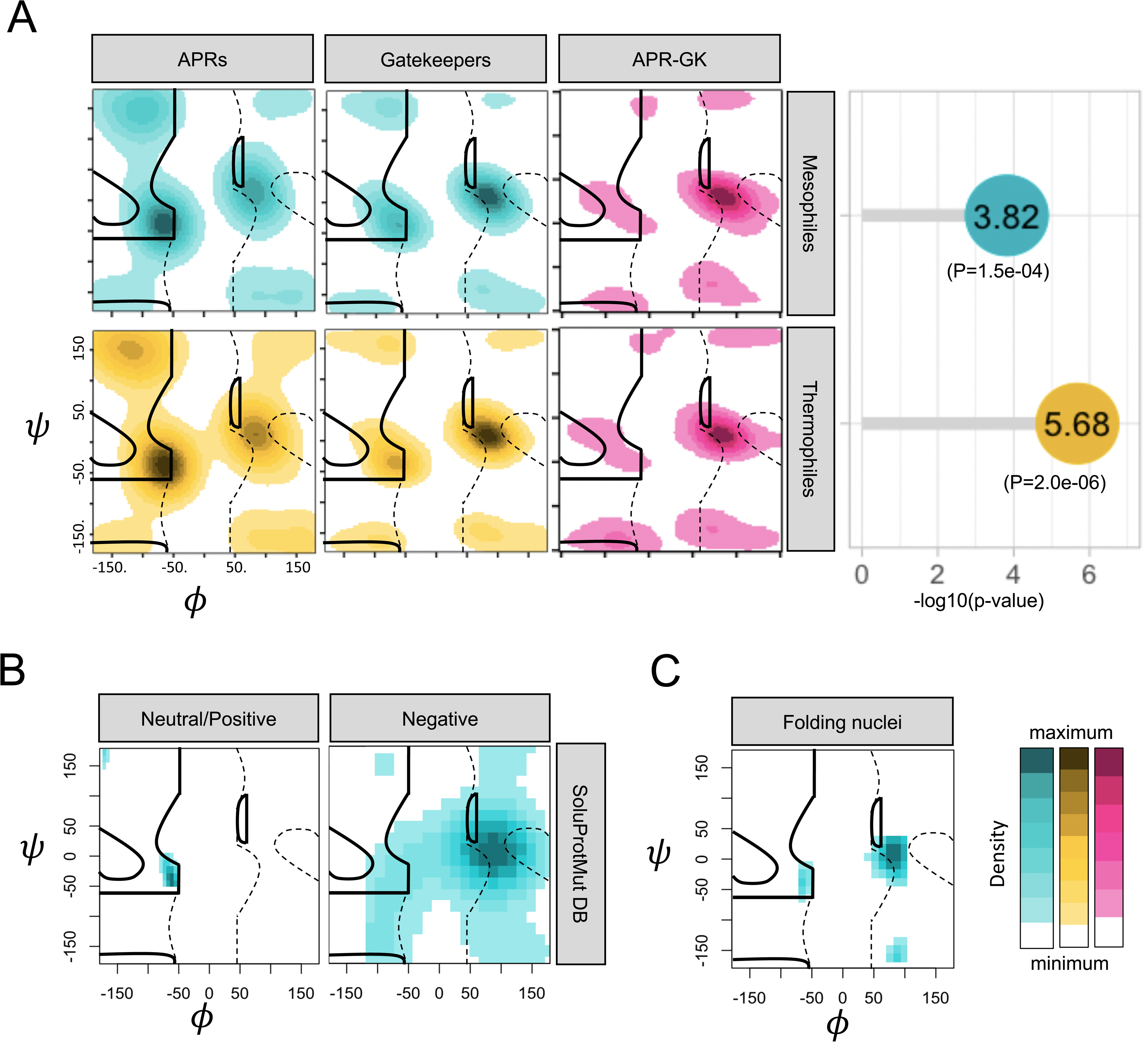
Conformation of glycyl residues present in aggregation-prone regions (APRs) and in flanking gate-keepers (GKs). (A) Ramachandran plots of glycyls in APRs and their flanking GKs (Thangakani et.al.) from mesophiles and thermophiles. The lollipop plot on the right signifies the P-values (-log10) of Chi-squared tests for the negative and positive phi values in APRs and GKs. (B) Ramachandran plots of glycyls that effects solubility positively or negatively or neutrally upon mutations to L-amino acids (SoluProtMut database) (C) Ramachandran plots of glycyls exhibiting high phi values of protein folding.

Since the protein stability and aggregation are inversely related to each other, we further tested the conformations of glycyl residues that exhibit higher ***Φ*** values (>0.4) of protein folding. We observed that the glycyl residues that were likely part of folding nucleus (high Φ values) preferably exhibited L-disallowed conformation (Figure 4C, Table S10). These observations coherently support that L-disallowed Gly are selected at sites that had role in promoting native folding and effectively avoiding inter-molecular aggregations.

### The plausible underlying mechanism

What might explain the observed association of L-disallowed Gly with the conformational stability and avoidance of aggregation? To address this, we first counted the L-disallowed glycyl residues that were associated with β-strands. Around 17% of all β-strand associated L-disallowed Gly were part of turns including β-turns and β-hairpins, the known motifs containing left-handed glycyl residues (Figure 5A). A majority (68%) of L-disallowed glycyls were close to the terminals of β -strands, while the rest 14% were interior to β-strand (Figure 5A). A closer inspection of the cases having L-disallowed Gly adjacent to or within the β-strand revealed that the L-disallowed Gly interrupts the alternating β-pleats by flipping or disorienting its Cα, interrupting the alternating phases of the β-pleats by incorrectly placing two or more consecutive residues towards one side (Figure 5B, Figure S4). This may or may not alter the atypical hydrogen bonding pattern of the β-sheet (Figure 4B). The disruption of the standard hydrogen bond pattern may imply β-bulges^29,30^, but not the lack thereof. We, therefore, referred these structures jointly as ‘β-crescents’ for the sake of convenience. The β-crescents can be approximated by greater distance between the Cα atoms of the preceding and proceeding residue of Gly (Table S11). Using this proxy, we showed that L-disallowed Gly tends to make crescent within or towards the end of the β-strand when compared with the L-allowed Gly and other residues in the sheet (Figure 5C). Importantly, regardless of hydrogen bond pattern, the crescents may disfavour extension of β-sheets. This is illustrated in the example given in Figure 5B. The ribbon model illustrates that the forthcoming β-strand on the left is interrupted just next to the Gly215 position (Figure 5B, lower panel).

**Figure 5.**
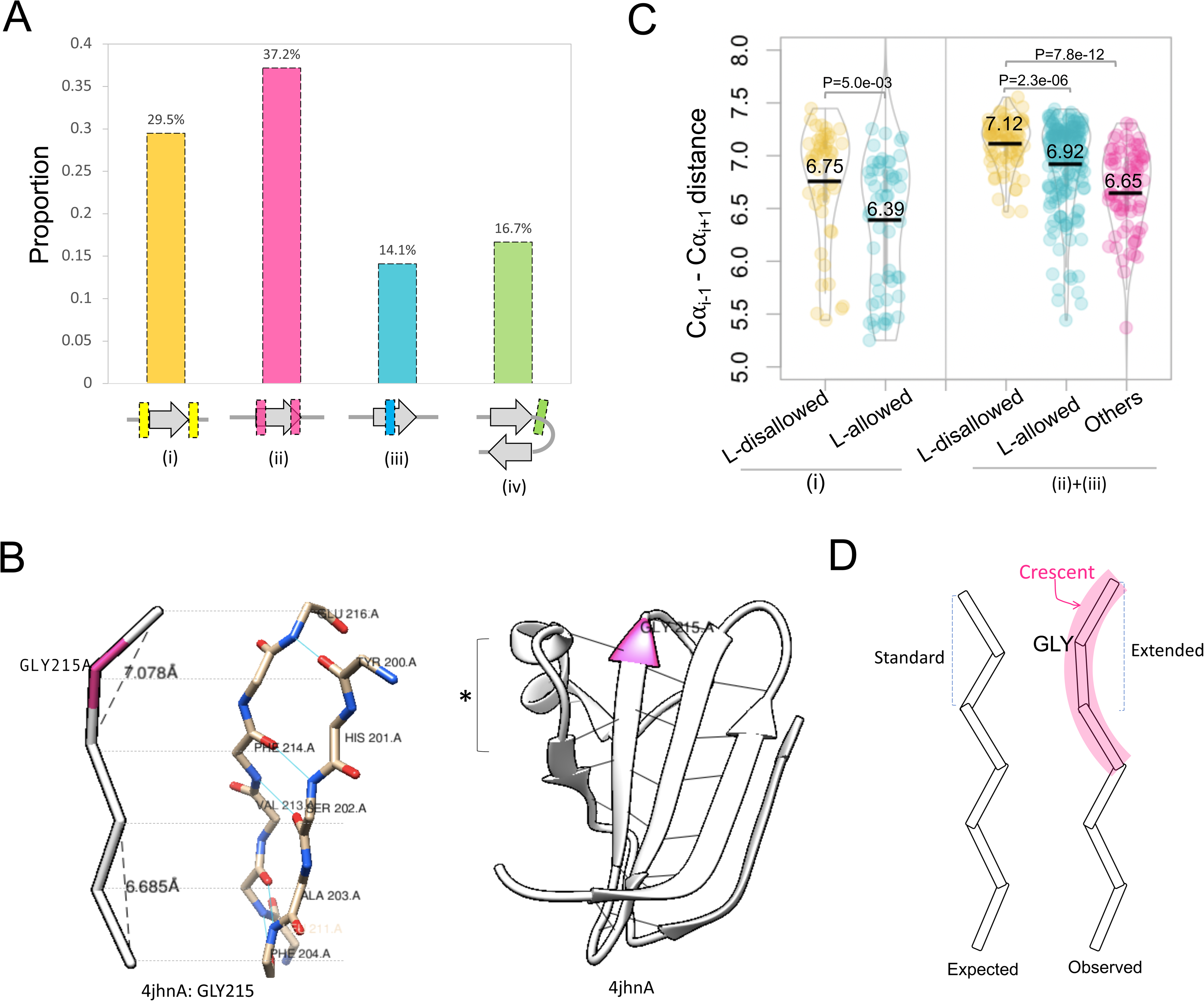
L-disallowed glycyl residues within or flanking β-strands. (A) Proportion of different scenarios of L-disallowed Gly associated with β-strands. L-disallowed Gly (i) immediately flanking β-strand;(ii) located at the terminal of β-strands; (iii) within β-strands, (iv) in turns. (B) Example illustrating β-crescent. Shown is the Cα trace, complete backbone, and the ribbon representations. The C***α*** of Gly215 should normally be facing inwards but has flipped due to positive " torsion. Vice-versa, the residue proceeding to Gly orients inwards instead of outwards. The vicinity of Gly215 does not show alteration of the Hydrogen bond on the right side but discourages the β-sheet expansion on left (marked as asterisk). (C) Distribution of C⍺_i-1_ to C⍺_i+1_ distances, where *i* marks the position of Gly, for the (i) L-disallowed and L-allowed Gly that were immediately flanking the β-strand, (ii) ones present towards terminals, (iii) ones within β-strands. We calculated p-values using Mann-Whitney U tests. (D) Model representation of β-crescent.

Parrini et al.^31^ has earlier shown that the glycyl residues are conserved in Acylphosphatase due to its role in avoiding aggregation. We obtained the glycyl residues associated with aggregation avoidance from their study and tested their conformation. These Gly consistently showed L-disallowed conformation (Figure 6A, Table S12). Interestingly, not only the sequence, L-disallowed conformations of Gly also had deep conservation across species. Out of total 6 such glycyls, 3 were located within or at the terminal of the β-strand. Gly15 was involved in a β-bulge, while Gly37 and Gly54 formed ‘β-crescents’ without alteration in the hydrogen bond pattern of β-strands (Figure 6B). The example illustrated that the conformational preferences of the conserved glycyl residues had been overlooked in the past.

**Figure 6.**
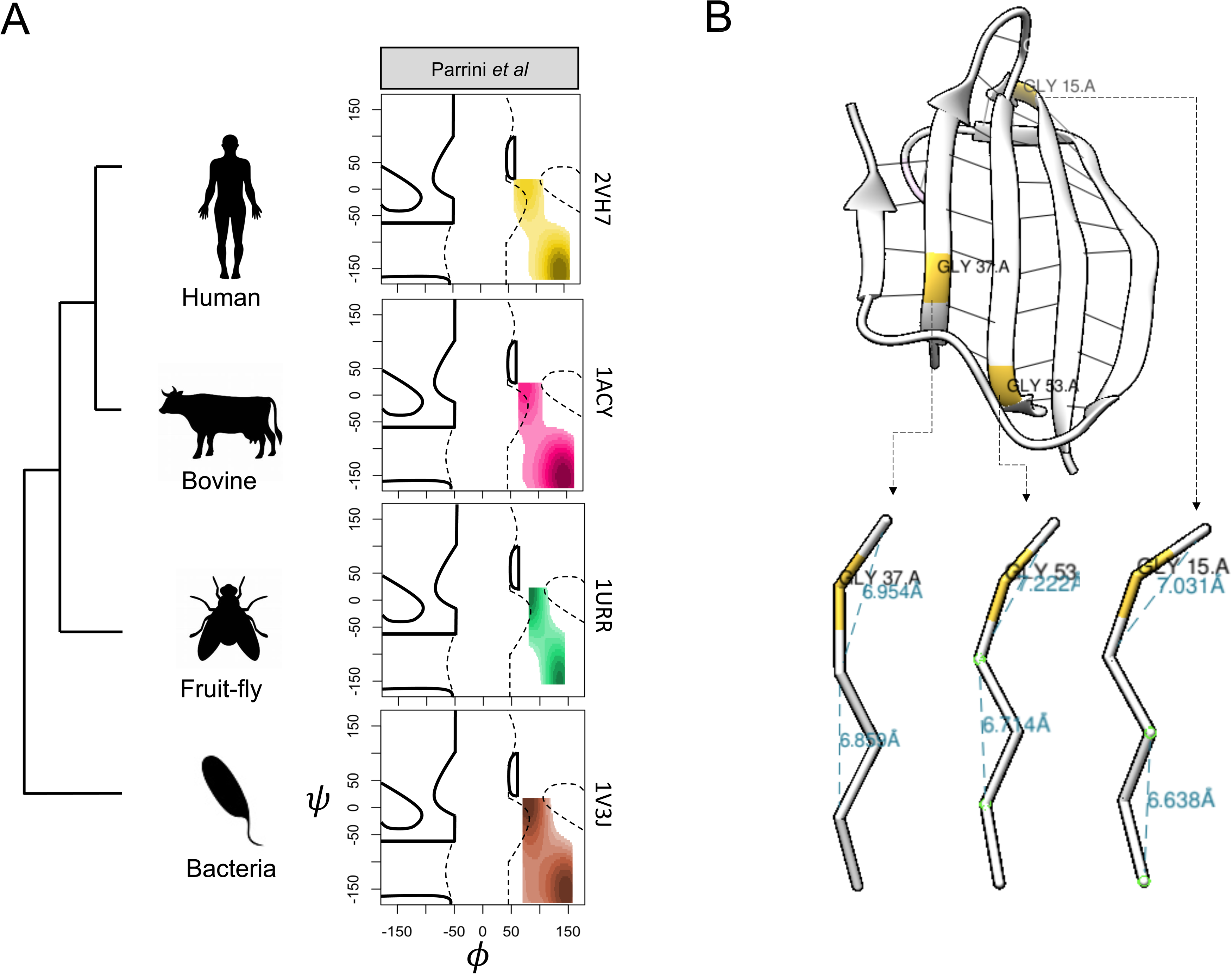
Conformation of conserved glycyl residues in Acylphosphatase (AcP). (A) Ramachandran plots of conserved glycyl residues in AcPs from phylogenetically distant species. (B) Locations and conformations of three β-crescents in AcP.

### L-disallowed conformation predicts the glycyl residues having pathogenic variants

We further asked if our observations had predictive power to prioritize the glycyl residues implicated in diseases. We implemented a deep learning method, namely Convolutional Neural Network (CNN), and showed that the model’s accuracy increased by 11% merely by adding glycyl conformation information in binary (1 for L-disallowed and 0 for L-allowed) (Figure 7A). Accordingly, the AUC of ROC curves improves by 9% (Figure 7B). We observed a similar trend for the XgBoost model. Figure S5 showed that the XgBoost model indeed used glycyl conformation in making decision through its boosted trees to infer pathogenic variants. The accuracy of our model increased dramatically to 93%, with a 4% improvement by adding glycyl conformation, when added the sequence information of adjacent regions of the glycyl residues (Figure S6). These results implied that the glycyl conformation exhibits predictive power to help prioritize pathogenic variants.

**Figure 7.**
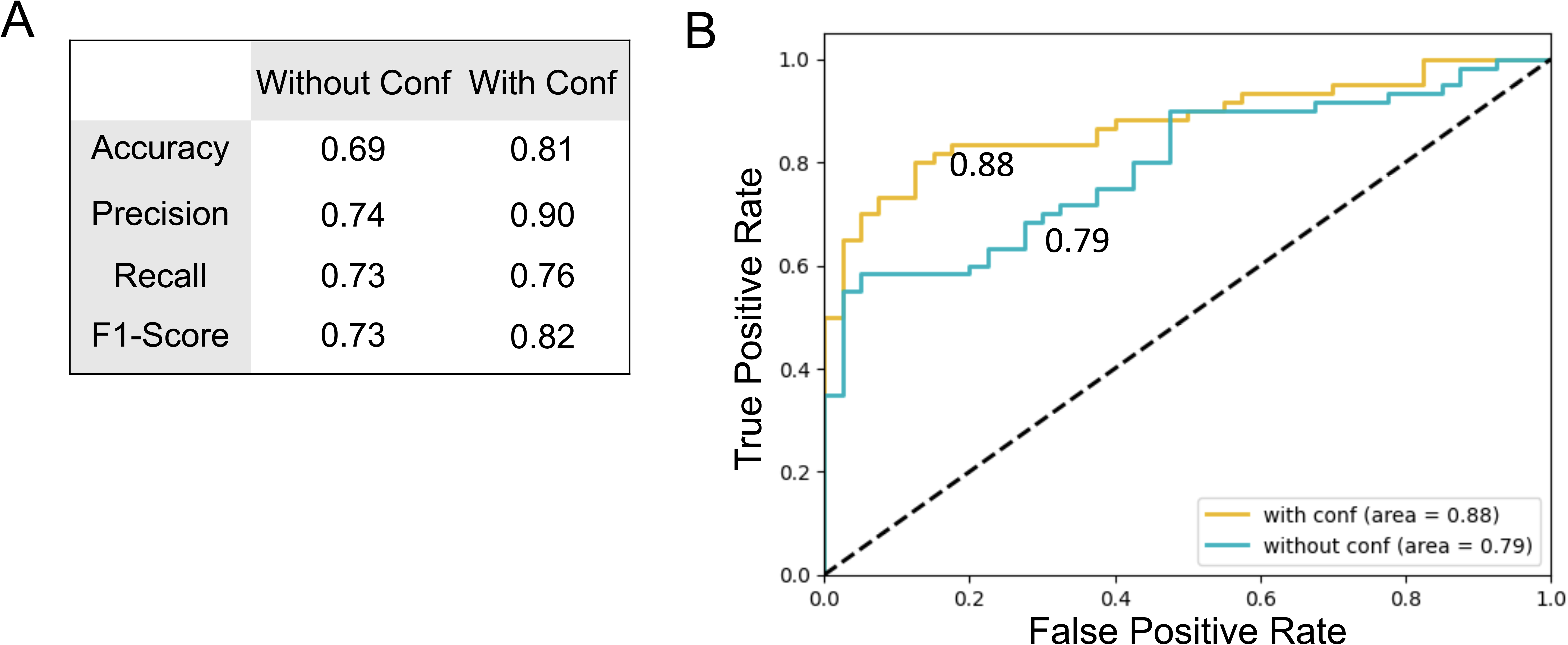
Predictive power of L-disallowed glycyls. (A) ROC curves (true positive rate as a function of false positive rate) for prediction of disease-associated glycyls with (gold) and without (aquamarine) considering glycyl conformation minimally (disallowed:1 and L-allowed:0) in the convolutional neural network (CNN) model. The numbers on the curves indicate the area under the curves (AUCs). (B) Performance measures of CNN model with and without glycyl conformation.

## Discussion

It remains challenging to identify the disease associations of missense variants since only a small subset of all missense variants link to pathogenicity. A large number of tools and algorithms have been developed to address this challenge^5–15^. Most of the algorithms rely on evolutionary information, surface accessibility, secondary structure, localization within functional sites, overlap with post-translational modification sites, type of amino acid, impact on thermostability, etc^5–15^. Little has been done to prioritize the residues based on conformational preferences. In this study, we demonstrated that the conformation of glycyl residues is critical to their association with pathogenicity. In particular, L-disallowed glycyls are relatively intolerable to missense mutations when compared to L-allowed ones. Higher conservation, the adverse effect on the protein stability upon mutation, enrichment near β-strand and within aggregation gatekeepers is the dominant pattern observed for the L-disallowed Gly at pathogenic sites. How does L-disallowed Gly help in stabilizing conformation and avoiding aggregation? The visualization of structures suggested that L-disallowed glycyl conforms a structural motif, referred here as β-crescents, wherein the alternation of β-pleats is interrupted and two or more residues including L-disallowed gly faces the same side of the β-strand forming a crescent like structure. These structures are akin to β-bulges though do not always interrupt the hydrogen bonding pattern of β-sheet. Interestingly, 29% of all structures were present at the edge strand. These cases may imply an evolutionary strategy to safeguard the native structure from inter-molecular aggregation. Indeed, β-bulges are earlier hypothesized to have a role in aggregation escape^32^. Nevertheless, the majority cases involve internal β-strands and may thus need other explanations. The energy barrier to form β-crescent would be greater for L-amino acids since ***Φ****-****Ψ*** dihedral angles for these L-amino acids cannot occupy L-disallowed space in Ramachandran plot without any steric clash. Modelling of mutations suggested that 70% of all Gly-to-Ala mutations conforms to D-conformers (positive, L-disallowed Φ values), implying the energetically incompatible conformation of L-Ala for L-disallowed glycyl residues (Table S13). Indeed, mutation of a conserved L-disallowed Gly to L-Ala leads to loss of proper folding and gain of inter-molecular aggregation of mammalian defensin proteins^33,34^, while mutation to D-Ala restores the proper folding of the β-bulge^35^. Further, the mutations of L-disallowed Gly to L-allowed amino acid may lead to correction of the Cα orientations to follow the proper alternation of β-pleats, and hence allowing the inter-molecular aggregation of folding intermediates. Since the positively charged gatekeeper residues (R and K) are the common targets of molecular chaperones, it is also likely that the L-disallowed glycyls, which are often found adjacent to R, could serve as chaperone target sites. Therefore, the L-disallowed conformation of Gly might be a prerequisite for the stabilization of folding intermediates. This was also supported by our observations that the glycyls present in folding nuclei tend to be L-disallowed. The mutation thereof may weaken the energy barrier between the native folded state and the aggregated state, i.e., shifting the kinetics towards aggregation, during the folding process.

Relatively weaker significance for the buried residues can be understood by the fact that the mutations of buried residues tend be destabilizing than those of surface. Greater enrichment of buried residues in the disease data may, therefore, blur the difference between the two alternate conformations. The surface residues might also be under greater evolutionary constraint to avoid the inter-molecular edge-to-edge β-strand aggregation among soluble proteins.

## Conclusion

Our findings strongly support that the mirror conformation of glycyl residues might have been evolutionarily selected at certain sites to promote native folding and avoid inter-molecular aggregation of proteins. Our observations had predictive power to help prioritize the sites with pathogenic missense variants and design proteins or peptides with improved stability and solubility.

## Method

### Pathogenic variants of Glycyl residues

We obtained the glycyl mutations associated with diseased or benign phenotypes from the ADDRESS (https://seq2fun.dcmb.med.umich.edu/ADDRESS/byStruct001narrow.html) database^36^.

### Sequence attributes

We fetched PDB’s SEQRES entries to calculate the amnio acid composition up- and - downstream of glycyl residues. The mean representation of 20 amino acids were calculated, and the pathogenic-to-benign ratio of those mean values were used for plotting purpose. We obtained the residue-wise evolutionary rates from ConSurfDB^37^ (https://consurfdb.tau.ac.il) to assess the conservation of L-allowed and disallowed glycyl residues.

### Structural attributes

The protein structures were fetched from RCSB PDB^38^ (https://www.rcsb.org/PDB). We wrote an in-house script for batch download. We mapped the residue-wise secondary structure states, and accessible surface areas (ASA) data using DSSP utility^39^ (https://swift.cmbi.umcn.nl/gv/dssp/), which was automated for bulk execution. Secondary structure states were further compressed into 4 broad states: ⍺-helix (H), extended β-sheet(E), turns(T) and unassigned(U). Secondary structure states for +/- 10 residues from the Gly were obtained and mean enrichments of different states were plotted. The turns were arbitrarily identified by considering the separation of two β-strands by 2-5 residues in T or U state in between. For surface accessibility, we considered a residue to be surface exposed if its ASA was at least 25% of the residue’s standard surface area. The calculation of dihedral angles and L-allowed/disallowed assignment was done using in-house scripts. We considered the following "- # ranges to demarcate the boundaries of L-allowed space in Ramachandran plot: (": 0° to -30°, #: -80° to 180°), (": 0° to -30°, #: -180° to -150°), and (": -30° to 90°, #: -10° to 120°). These ranges had been earlier used by others for the same purpose^40^. We measured C⍺_i-1_ to C⍺_i+1_ distances using inhouse script. VMD^41^ (https://www.ks.uiuc.edu/Research/vmd/) and Chimera^42^ (https://www.cgl.ucsf.edu/chimera/) were used for structure visualizations. We obtained the β-bridge information from DSSP files to assess the symmetry of H-bonds around β-strands. We designated a β-strand as edge strand if five consecutive residues had H-bonds only on one side of the β-strand. All H-bonds formed by the glycyl residues were obtained using the HBOND program (http://cib.cf.ocha.ac.jp/bitool/HBOND/) looped for batch run.

### Protein stability

We obtained the change in Gibbs free energy upon mutation (ΔΔG) by fetching FoldX entries in the ADDRESS database. The experimental ΔΔG were obtained from the ProTherm database^43^(https://web.iitm.ac.in/bioinfo2/prothermdb/index.html). The following equation was considered: ΔΔG = ΔG_variant_ - ΔG_WT_. Since FoldX provides the ΔΔG values using ΔG_WT_ - ΔG_variant_ expression, we inversed the sign for FoldX’s ΔΔG values to maintain the concordance with ProTherm.

### Protein aggregation and solubility

We used the AGGRESCAN^44^ (http://bioinf.uab.es/aggrescan/) package to calculate aggregation propensity (*‘a4v’* values) for the regions up- and downstream to the glycyl residues. The aggregation potential values were obtained for +/- 10 residues from Gly and the mean for each position was calculated. We obtained experimentally measured solubility changes upon mutations from manually curated SoluProtMut database^28^ (http://bioinf.uab.es/aggrescan/).The folding nucleus data (">0.4) was obtained from literature^45-47^ (http://bioinfo.protres.ru/foldnucleus/). The APR and GK data for 373 thermophilic and 373 mesophilic proteins was obtained from Thangakani et.al.^27^.

### Mutation modelling

We modelled Gly-to-Ala mutations using Modeller^46^ (https://salilab.org/modeller/download_installation.html).

### Statistics

Odd ratios were calculated using the following equation:

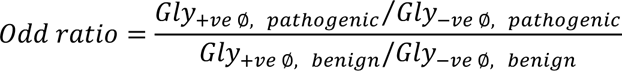

Lower 95% confidence intervals were calculated using the following equation:

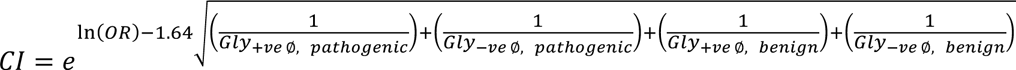

We performed Fisher’s exact tests and Chi-squared tests using ‘fisher.test’ and ‘chisq.test’ functions in R. Pearson’s residuals were calculated using the following equation:

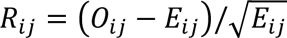

Where O_ij_ and E_ij_ observed and expected counts respectively in ith column and jth row. The Pearson’s residuals signify the positive or negative associations between column and row entities of the contingency table. The balloon-plots were made using ‘gplots’ R-package(https://cran.r-project.org/web/packages/gplots/). Mann-Whitney U and t-tests were performed using ‘wilcox.test’ and ‘t.test’ functions respectively in R. We used ‘ggplot2’ R-package to make multi-facet boxplots (https://cran.r-project.org/web/packages/ggplot2/). Ramachandran plots were drawn using 2D kernel density function ‘kde2d’ of ‘MASS’ R-package(https://cran.r-project.org/web/packages/MASS/).

### Machine Learning

We implemented ‘tensorflow’ (https://pypi.org/project/tensorflow/) and ‘xgboost’ (https://pypi.org/project/xgboost/) python packages for CNN and XgBoost models. Glycyl conformation (L-allowed:0, L-disallowed:1), FoldX’s ΔΔG, number of short contacts, number of H-bonds, residue-wise evolutionary rates, and ASA values were considered as training attributes for pathogenic and benign variants of glycyl residues. Python codes are shared at: https://github.com/rajeshpotlia/glycyl-ml.

## Acknowledgement

The authors duly acknowledge the financial support from Department of Biotechnology, India (BT/PR40149/BTIS/137/36/2022, BT/PR40198/BTIS/137/56/2023).

## Supplementary Figures Legends

**Figure S1.**
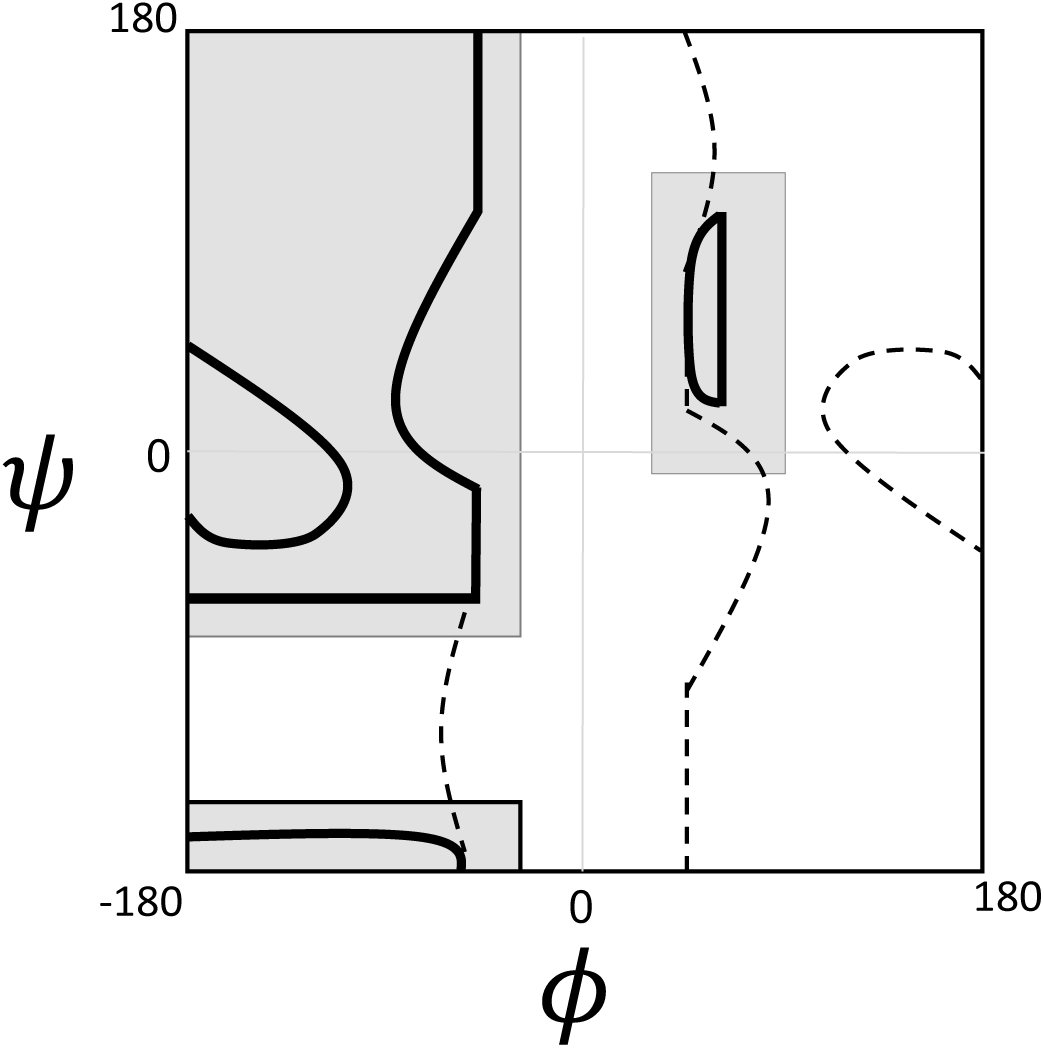
Schematic representation of boundaries to determine the L-allowed (grey shaded area) and L-disallowed (white area enclosed by dotted lines) conformation of glycyl residues. Top left: (": -180° to -30°, #: -80° to 180°), bottom left: (": -180° to -30°, #: -180° to -150°), and right middle: (": 30° to 90°, #: -10° to 120°)

**Figure S2.**
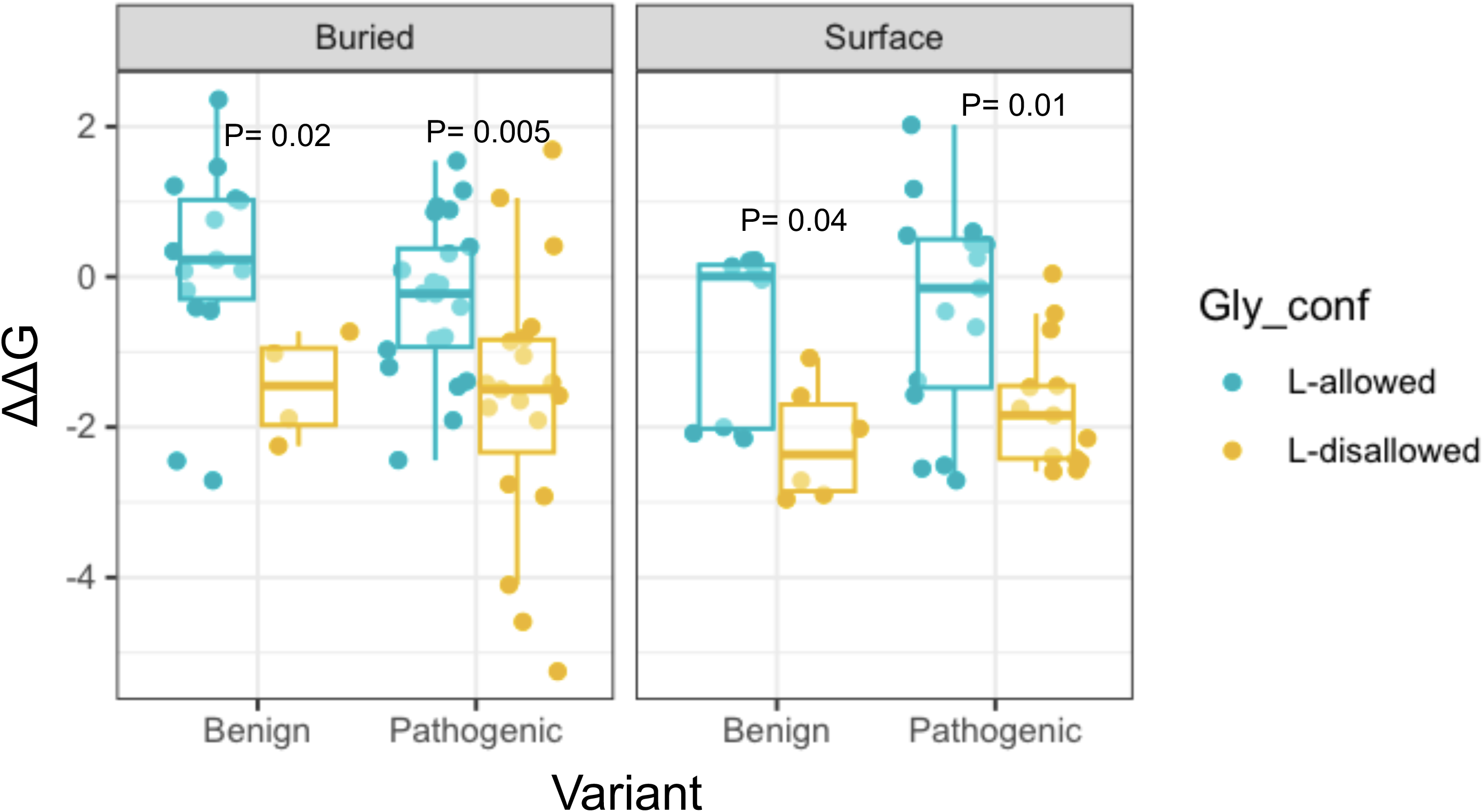
Gibbs free energy changes (upon mutation) for L-allowed and L-disallowed glycyl residues that were mutated to Ala.

**Figure S3.**
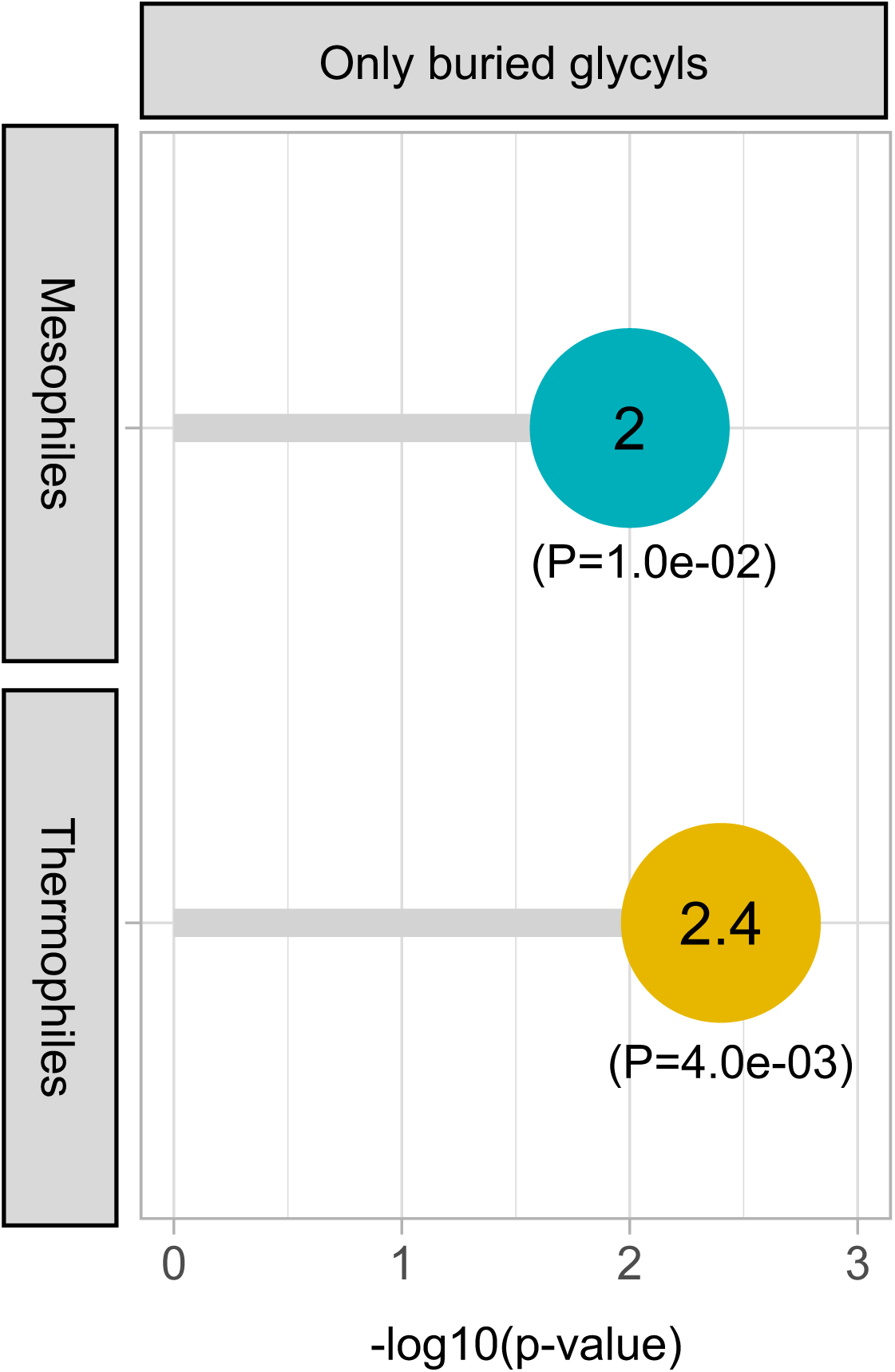
The P-values (-log10) of Chi-squared tests for the negative and positive phi values of buried residues in APRs and GKs.

**Figure S4.**
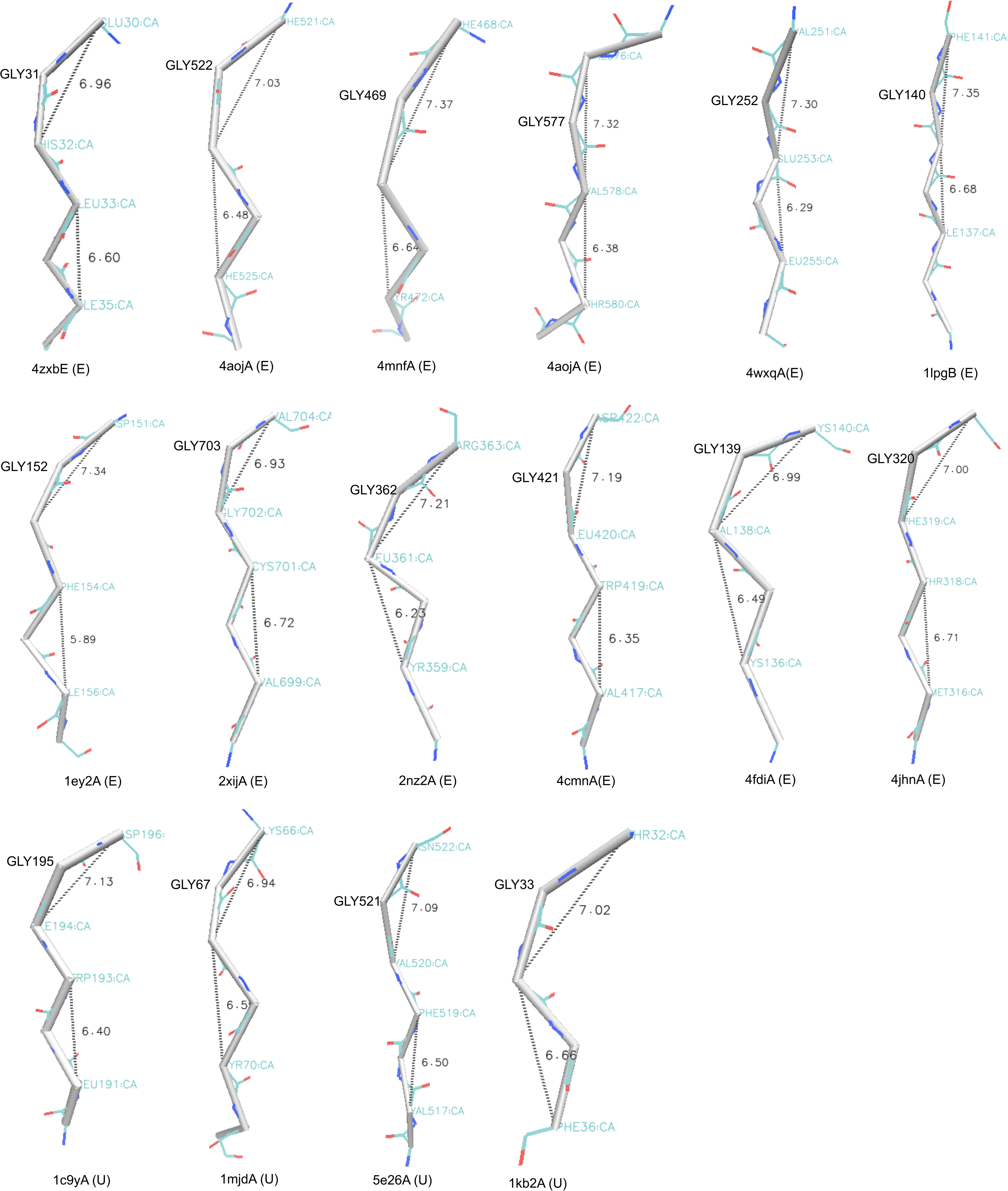
Visual examples of β-crescents. CA-traces superimposed onto the backbone were drawn using VMD.

**Figure S5.**
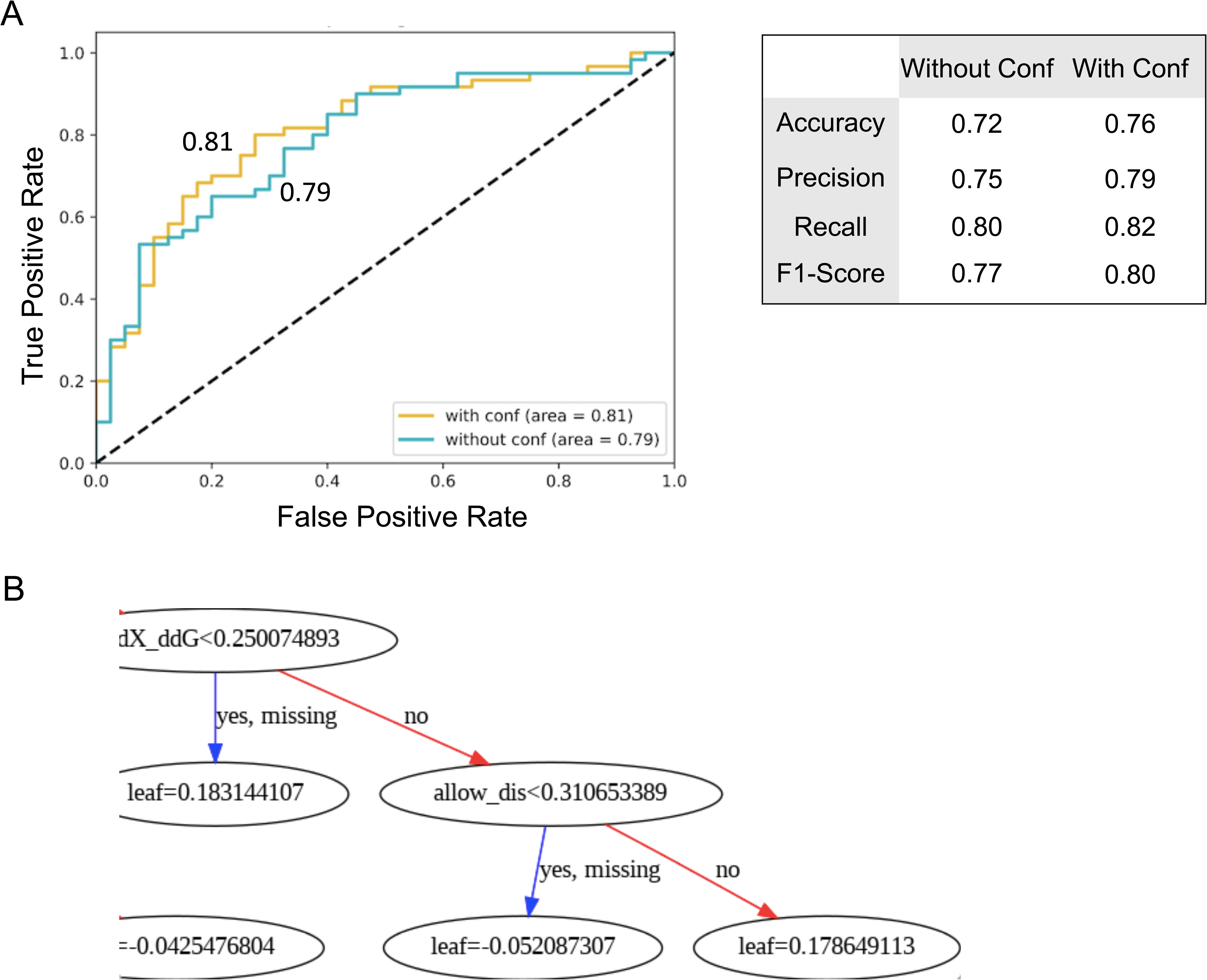
Performance of XgBoost model for the prediction of disease-associated glycyls. (A) ROC curves with (gold) and without (aquamarine) including glycyl conformations among training attributes. The numbers on the curves indicate the area under the curves (AUCs). (B) Performance measures of XgBoost model with and without including glycyl conformation. (C) An example of boosted tree illustrating that the model uses glycyl conformation in making decisions.

**Figure S6.**
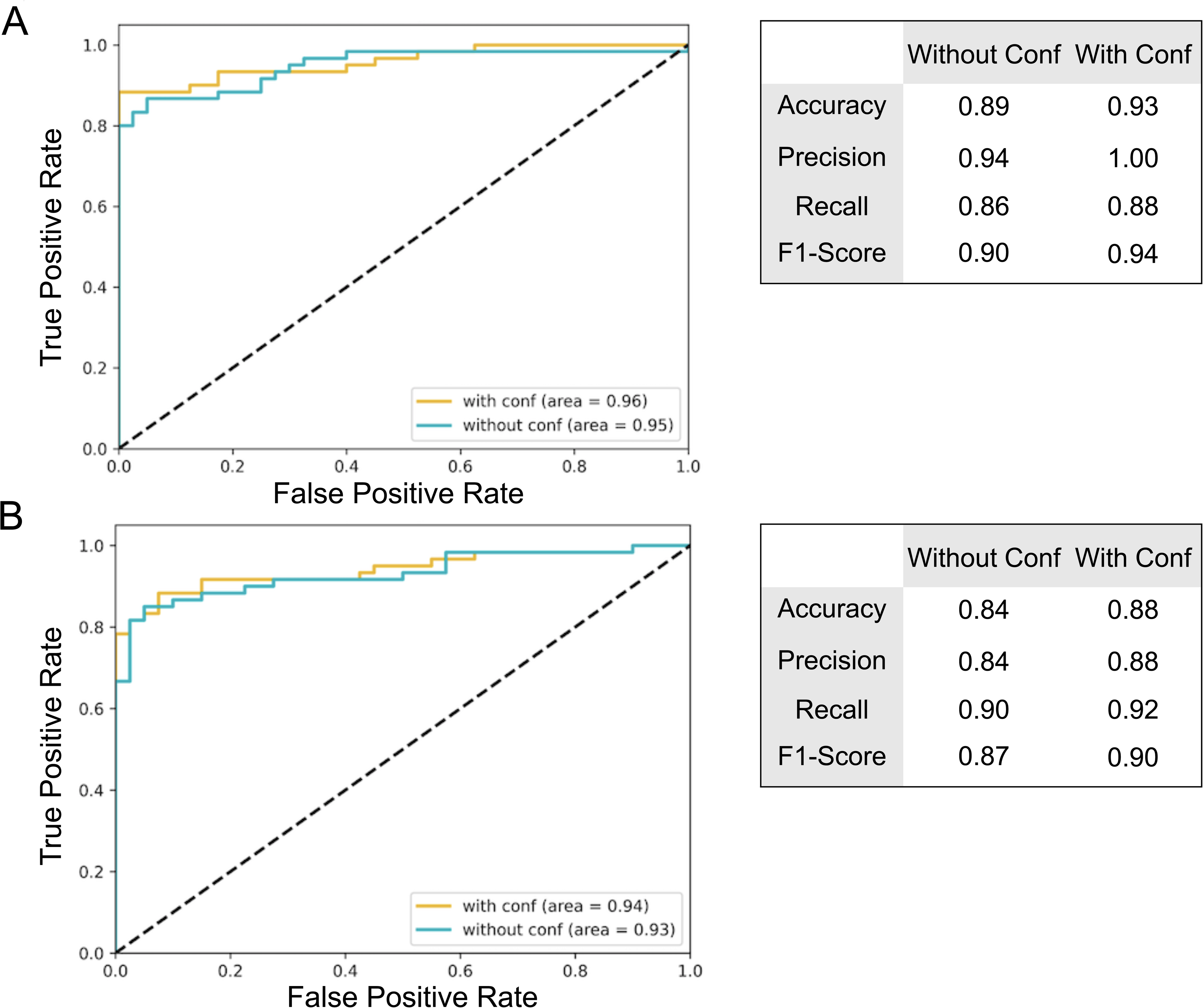
Predictive power of glycyl conformation when added the amino acid composition of adjacent regions. (A) ROC curves (true positive rate as a function of false positive rate) for prediction of disease-associated glycyls with (gold) and without (aquamarine) considering glycyl conformation minimally (disallowed:1 and L-allowed:0) in the convolutional neural network (CNN) model. The numbers on the curves indicate the area under the curves (AUCs). (B) Performance measures of CNN model with and without glycyl conformation. (C-D) Same as A-B, but for the XgBoost model.

## Supplementary Tables

**Table S1.** Dihedral angles, secondary structures, evolutionary divergence, and FoldX-inferred ΔΔG values of Glycyl residues having missense variants.

**Table S2.** Annotations of glycyl residues at pathogenic and benign sites through ccPDB.

**Table S3.** Dihedral angles and experimental ΔΔG values (from ProTherm database) of glycyl residues.

**Table S4.** Dihedral angles and FoldX ΔΔG values for the disease-associated glycyls that had alanine variants.

**Table S5.** Secondary structure states of +/- 10 residues to glycyls.

**Table S6.** Amino acid composition upstream and downstream to glycyl residues.

**Table S7.** AGGRESCAN scores around glycyl residues.

**Table S8.** Dihedral angles of glycyls present in APRs and GKs of 373 homologous mesophiles and corresponding thermophiles.

**Table S9.** Dihedral angles of glycyls exhibiting change in protein solubility upon mutation.

**Table S10.** Dihedral angles of glycyls present in the folding nuclei

**Table S11.** C⍺_i-1_ to C⍺_i+1_ distances for the glycyls present within or towards terminals (inside/outside) of b-strands.

**Table S12.** Dihedral angles of conserved glycyls in Acylphosphatase.

**Table S13.** Dihedral angles for the modelled Gly-to-Ala mutations for all L-disallowed glycyls present in ADDRESS database.

